# OCD.py - Characterizing immunoglobulin inter-domain orientations

**DOI:** 10.1101/2021.03.15.435379

**Authors:** Valentin J. Hoerschinger, Monica L. Fernández-Quintero, Franz Waibl, Johannes Kraml, Alexander Bujotzek, Guy Georges, Klaus R. Liedl

**Affiliations:** Department of General, Inorganic and Theoretical Chemistry, and Center for Molecular Biosciences Innsbruck (CMBI), University of Innsbruck, Innrain 80-82, A-6020 Innsbruck, Austria; Roche Pharma Research and Early Development, Large Molecule Research, Roche Innovation Center Munich, Penzberg, Germany

## Abstract

**Summary:** Inter-domain orientations between immunoglobulin domains are important for the modeling and engineering of novel antibody therapeutics. Previous tools to describe these orientations are applicable only to the variable domains of antibodies and T-cell receptors. We present the “Orientation of Cylindrical Domains (OCD)” tool, which employs a transferable approach to calculate inter-domain orientations for all immunoglobulin domains. Based on a reference structure, the OCD tool automatically builds a suitable reference coordinate system for each domain. Through alignment, the reference coordinate systems are transferred onto the sample to calculate six measures which fully characterize the inter-domain orientation.

**Availability and implementation:** The OCD approach is implemented as a stand-alone Python script, OCD.py, which can handle multiple types of data input for the analysis of single structures and molecular dynamics trajectories alike. OCD.py is available at https://github.com/liedllab/OCD under MIT license.

**Supplementary Information:** Supplementary information and data are available at *Bioinformatics* online.

## Introduction

Antibodies are one of the fastest growing classes of biotherapeutic proteins on the market, generating an estimated revenue of $115 billion in 2018 with an additional projected growth to $300 billion in 2025 (Lu *et al.*, 2020) and a significant amount of antibody biotherapeutics in the pipeline (Kaplon and Reichert, 2021). For modeling and engineering such antibodies, the relative interface orientation between two immunoglobulin domains plays a central role. For example, the inter-domain orientation of variable immunoglobulin domains in antibody and T-cell receptors has been linked to CDR loop conformations and therefore influences the paratope structure (Dunbar *et al.*, 2013; Bujotzek *et al.*, 2015; Fernández‐Quintero *et al.*, 2020). While computational tools to fully characterize the Fv region of antibodies and TCRs are already available (Dunbar *et al.*, 2013; Marze *et al.*, 2016), no such tools were published for other immunoglobulin domain interfaces, such as the C_H_3-C_H_3 and the C_H_1-C_L_ interface.

We therefore present the “Orientation of Cylindrical Domains (OCD)” tool, an easily transferable approach to describe inter-domain orientations for a variety of protein domains. The Python-based tool, OCD.py, can handle PDB crystal structures and molecular dynamics (MD) simulation trajectories files. The script uses the pytraj (Nguyen *et al.*, 2016), mdtraj (McGibbon *et al.*, 2015), pandas (McKinney, 2010) and NumPy (Harris *et al.*, 2020) libraries for rapid handling of structural data, structure alignment and internal calculations. For a list of supported filetypes, please refer to pytraj’s documentation. The tool plots the generated orientational data using Matplotlibs’ pyplot library (Hunter, 2007) and provides visualizations in the form of VMD (Humphrey *et al.*, 1996) and PyMol (Schrödinger LLC, 2020) input scripts. The calculation time of a C_H_3-C_H_3 interface is found to be less than a second for a single PDB crystal structure and under a minute for a 50.000 frame trajectory (see SI section 2). The OCD tool is therefore well suited as a standard analysis tool for MD trajectories of immunoglobulin domains. OCD.py is available at https://github.com/liedllab/OCD. Example reference structures and atom masks are provided for the IgG1 Fv and C_H_3 regions.

## OCD.py

Existing approaches, like the current state-of-the-art ABangle (Dunbar *et al.*, 2013), rely on pre-defined residues linked to inter-domain orientations and large amounts of structural data to define a coordinate system for the calculation of inter-domain orientations. The OCD approach expands on this by automatically creating a suitable coordinate system for the characterization of these interface orientations for any user-provided reference structure. This allows a straight-forward analysis without the significant demands on previous structural knowledge. Using this tool, a reference coordinate system is created based on user-defined reference structures consisting of an atomic structure and two domain selections over these atoms. To this end, the reference structure for each domain is generated by considering a center axis linking the two centers of mass of the different domains, and the first principal axis P of inertia of each domain corresponding to the lowest eigenvalue of the inertia tensor. Each individual domain is aligned to the world coordinate system by aligning this principal axis to the z unit vector and the center axis as close as possible to the x unit vector, yielding a reference structure for each domain. To map the coordinate system onto a sample structure, the references are aligned to the sample and the alignment transformations are applied to the xyz unit vectors. The transformed z vectors (A1/ B1) and y vectors (A2 / B2) as well as the center axis are then used to calculate six orientational measures: Two tilt angles for each vector towards the center axis (AC1, AC2, BC1, BC2), the length of the center axis (dC) and a torsion angle (AB) between the two intersecting planes composed of A1, the center axis and B1. The generation of reference structures and coordinate system is outlined in figure 1.

**Figure 1.**
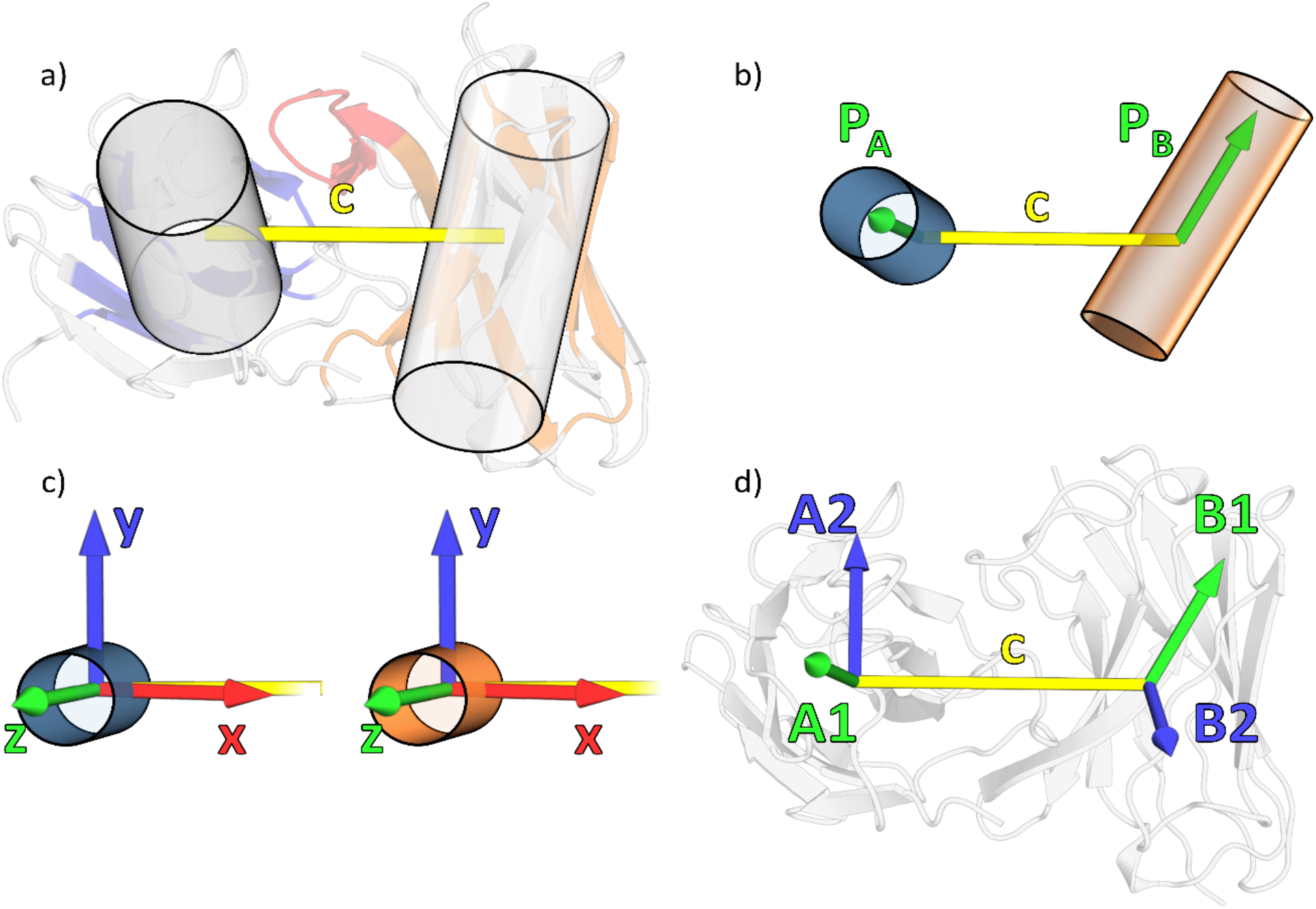
a) A user-supplied structure and two atom selections are used to generate two reference structures. b) After applying the atom selections, the principal axis **P** corresponding to the lowest eigenvalue is calculated for both domains. c) The domains are reoriented into a standard orientation, where **P** is aligned to the z unit vector and the center axis is as close as possible to the x unit vector. d) By aligning the reference structures to a sample and transferring the alignment to the z and y unit vectors, the OCD coordinate system is mapped onto the sample.

The intermediate protein structures for the generation of a V_H_ reference structure are shown in SI section 1.

The approach is transferable to different domain types, if a reasonable reference structure can be provided. There are two constraints necessary to generate a suitable coordinate system: (1) The reference structure needs to align well to all sample structures and (2) the first principal axis of both domains should not be parallel to the axis linking the domains’ centers of mass. Both criteria can be examined through the data and visualization output by the tool, enabling the user to easily verify their approach.

When using the same reference structures as ABangle, the V_H_−V_L_ orientations calculated by the OCD tool can be used to perfectly predict orientations obtained with ABangle through a multivariate linear regression (R^2^ = 0.99, see SI section 3). When using a reference structure created by averaging the Fv structures found in the ABangle dataset, we still find a high correlation (R^2^ = 0.85), thereby showing the OCD approach’s flexibility. As examples, we applied the OCD tool to the dataset of non-redundant Fv structures provided with ABangle as well as to C_H_3−C_H_3 MD simulations, which is documented in the SI section 4.

## Conclusion

The OCD tool extends previous approaches for the calculation of antibody interface orientations by involving the user in the choice of reference structure. It is transferable to a variety of protein domain interfaces if the outlined constraints are met. It is especially suited for the description of immunoglobulin domains. Additionally, it natively supports the fast analysis of MD trajectories, allowing for insights into static structural data as well as dynamics of the sampled interfaces.

