## Supporting Information for "OCD.py - Characterizing immunoglobulin inter-domain orientations"

### 1. OCD coordinate system for a $V_H$ domain

The process of generating the OCD coordinate system for a Fv system is shown in figure S1. Note, that while the process is here shown for only the VH domain, the same procedure is of course applied to the $V_L$ domain for the OCD calculation. First, the structure containing the two domains as supplied by the user is read in and the atom selections are applied (Figure S1 a). For each domain, the principal axis I corresponding to the lowest moment of ineratia eigenvalue is calculated for the atoms selected by the user – in the example these are parts of the beta-strands of the $V_H$ domain (Figure S1 b). Both domains are reoriented into a standard orientation, where the principal axis I is aligned to the z unit vector, the center of mass to the origin and the center axis c is brought as close as possible to the unit vector x (Figure S1 c). The two domains in the standard orientation are the reference structures, which are aligned to the sample, thereby mapping the OCD coordinate system to the sample (Figure S1 d). From this coordinate system, the OCD angles can then be calculated

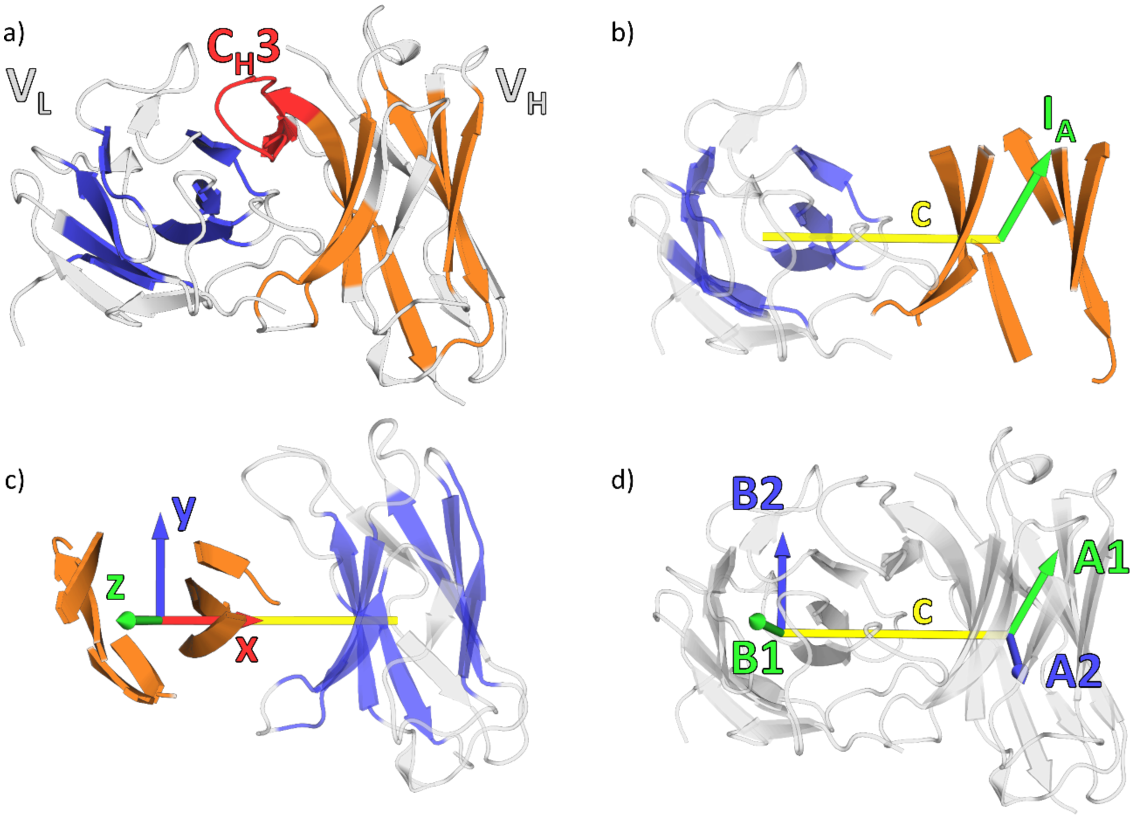


*Figure S 1: The process of generating the OCD coordinate system for a Fv system. a) User-supplied structure and atom selections (blue, orange). b) The principal axis I corresponding to the lowest eigenvalue is calculated for both atom selections (here only shown for $V_H$). c) The domains are then into a standard orientation, thereby forming the reference structures. D) The reference structures are aligned to the sample, inscribing the OCD coordinate system.*

### 2. Speed Benchmark

The calculation speed of OCD.py was benchmarked to give an outlook on its everyday usability. Both the performance on single pdb files and MD trajectories was measured. The benchmarks were run as single core calculations on an Intel i7-6800K CPU at 3.40 GHz.

For the pdb benchmark, structures from the ABangle dataset (Dunbar *et al.*, 2013) were analyzed. An increasing number of pdb files was calculated in each step and the total CPU calculation time recorded. For benchmarking purposes, the plotting features of the OCD tool were turned off. As expected, the calculation time increases linearly. By fitting the data with a linear regression, the calculation time per structure and the overhead are estimated. The data and the linear fit are shown in figure S1. The calculation time was found to be 0.73 s per pdb file with an additional overhead time of 0.12 s. Similarly, the benchmark for simulations was run using an increasing amount of frames of a MD simulation based on the $C_H3$-$C_H3$ crystal structure 3AVE (Matsumiya *et al.*, 2007) (for details see section "Applications to MD trajectories"). The data and linear fit are shown in figure S2.

The calculation time was found to be 0.0011 s per frame with 0.55 s of overhead. Clearly, analyzing simulation frames is far faster than single pdb files. This is most likely due to the internal file handling done by pytraj (Nguyen *et al.*, 2016), which enables the rapid analysis of trajectories in the OCD tool.

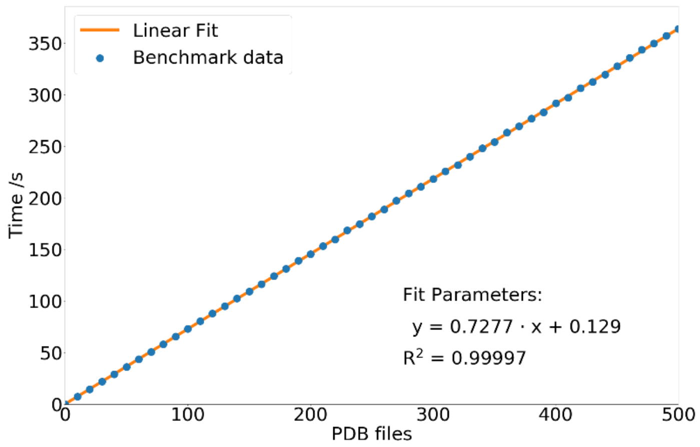


*Figure S 2: Benchmark for multiple runs using an increasing number of PDB files.*

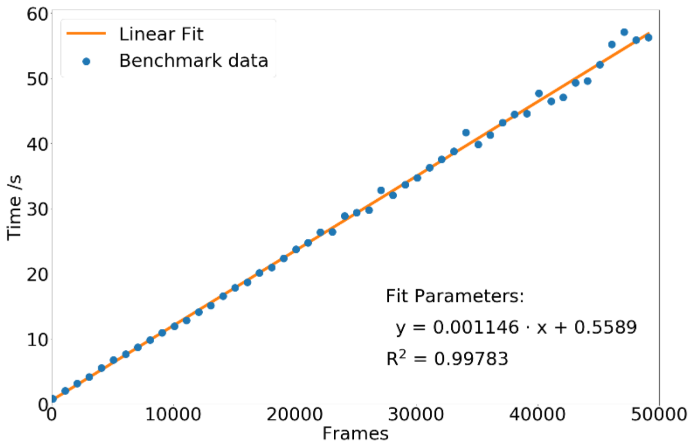


*Figure S 3: Benchmark for multiple runs using an increasing number of MD trajectory frames.*

### 3. Comparison with ABangle on the ABangle non-redundant dataset

To validate our approach, orientational measures of the non-redundant ABangle Fv dataset (Dunbar *et al.*, 2013) were calculated and compared to the ABangle results. To enable a direct comparison, the two ABangle reference structures were aligned onto the first entry in the ABangle Fv dataset, 1AOQ, and then combined into a single structure. Thereby, a full Fv reference structure was created which was then used to calculate the orientational measures using our OCD tool. Before the calculation, the CDR insertions (e.g. residues L30A, L30B, …) were removed from the structure files as OCD.py ignores insertion codes. As the choice of coordinate system is different between our approach and ABangle, a direct comparison of a single measure is not possible. We therefore tried to fit the data of our approach in a multivariate linear regression to the data obtained by ABangle, as shown in figure S2. In doing so, we obtained a coefficient of determination ($R^2$) of over 0.99. A 10-fold cross validation yielded an averaged $R^2$ of 0.9889 with a standard deviation of 0.006. The measures generated by ABangle therefore describe the same orientation as the measures calculated from our approach, though the measures themselves are different from each other.

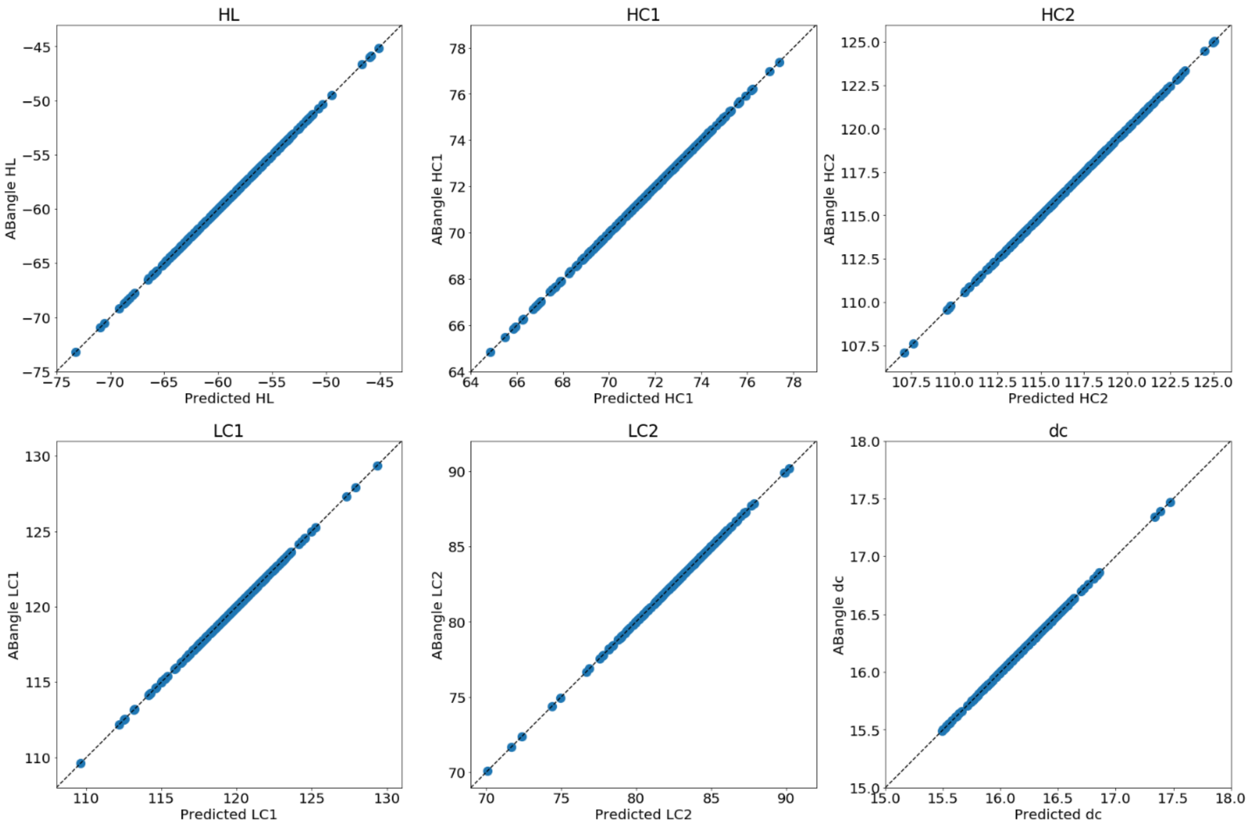


*Figure S 4: Measures calculated with ABangle compared to OCD measures fitted with a multivariant regression. The OCD calculation used the ABangle reference structure as reference. The diagonal is shown as a black dotted line.*

When a different Fv reference is used, the OCD orientation does not mirror the ABangle orientation as perfectly as before. We repeated the previous procedure but the reference structure this time was an averaged structure from all Fv pdb files found in the ABangle dataset. Additionally, the atom selection mask shown in figure S1 was applied. The results are shown in figure S5. For this fit, the overall coefficient of determination was 0.85. This decrease is most likely due to worse alignment and the difference in the analyzed orientation: while ABangle describes the orientation of the residues close to the interface, the OCD approach using the six beta-strands gives a broader overview of the overall domains towards each other. Additionally, three outliers (colored in red in the plots) with a high alignment RMSD of over 3 Å in the heavy chain were observed. After removal of the three outliers from the data, a 10-fold cross validation resulted in an averaged $R^2$ of 0.8662 with a standard deviation of 0.0300.

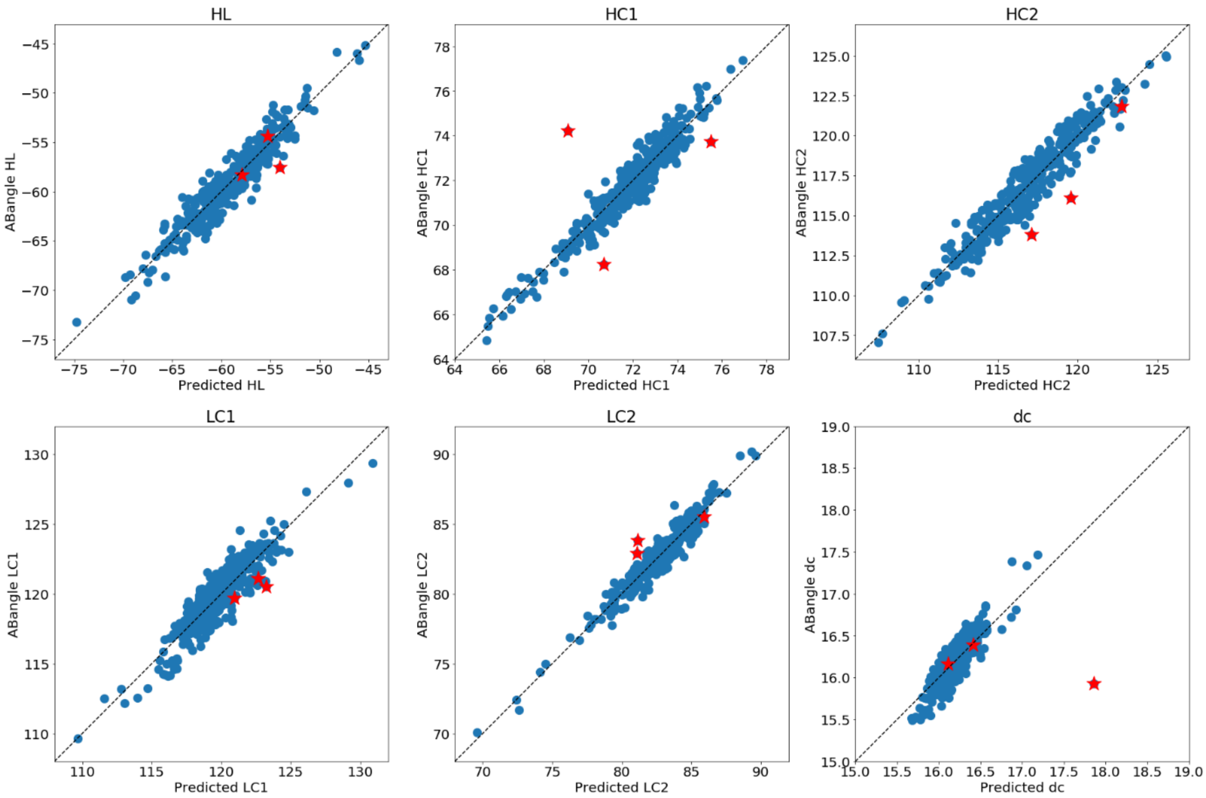


*Figure S 5: Measures calculated with ABangle compared to OCD measures fitted with a multivariant regression. The OCD calculation used an averaged Fv structure with an additional atom mask, restricting the used atoms to six beta-strands. The data shown as red stars represents three outliers with an alignment RMSD of over 3 Å. The diagonal is shown as a black dotted line.*

### 4. Application to MD trajectories

The OCD tool was applied to a simulation of a $C_H3$-$C_H3$ interface and a simulation of a $C_H1$-$C_L$ interface. To run these simulations, PDB structures 3AVE and 1N8Z were used as starting structures. The crystal structures were protonated using MOEs protonate3D function (Labute, 2009). 3AVE was cut to only include the $C_H3$, while the full Fab structure was used for 1N8Z. They were then solvated in a TIP3P cubic water box (Jorgensen *et al.*, 1983) with a minimum distance to the border of 10 Å for 1N8Z and 12 Å for 3AVE in AMBER's tLEaP (Case *et al.*, 2020). Uniform background charges were applied to neutralize residual charges in the system(Case *et al.*, 2020; Hub *et al.*, 2014). A coordinate and topology file of the structure using AMBERs ff14SB forcefield(Case *et al.*, 2020; Maier *et al.*, 2015) was saved and then equilibrated according to an extensive 20 step equilibration process (Wallnoefer *et al.*, 2011). Both simulations were run for 1 µs at a temperature of 300K and a pressure of 1 bar using pmemd.cuda (Salomon-Ferrer *et al.*, 2013; Le Grand *et al.*, 2013), a GPU accelerated version of AMBERs pmemd. Temperature and pressure were held constant with a Langevin thermostat (Loncharich *et al.*, 1992; Adelman and Doll, 1976) and a Berendsen barostat (Berendsen *et al.*, 1984) respectively. The Particle Mesh Ewald method (Darden *et al.*, 1993) was used to treat long range electrostatic interactions. Short range van der Waals interactions were cut off after 8 Å. Hydrogens were restricted according to the SHAKE algorithm, allowing for a time step of 2 fs (Miyamoto and Kollman, 1992).

The OCD measures calculated of the $C_H3$-$C_H3$ simulation are shown in figure S6, the results for the $C_H1$-$C_L$ simulation in figure S7. For the calculation of the $C_H3$-$C_H3$ simulation's measures, an atom mask encompassing only Cα atoms in the six major beta-strands was used. In the calculation of the $C_H1$-$C_L$ simulation all Cα atoms were used.

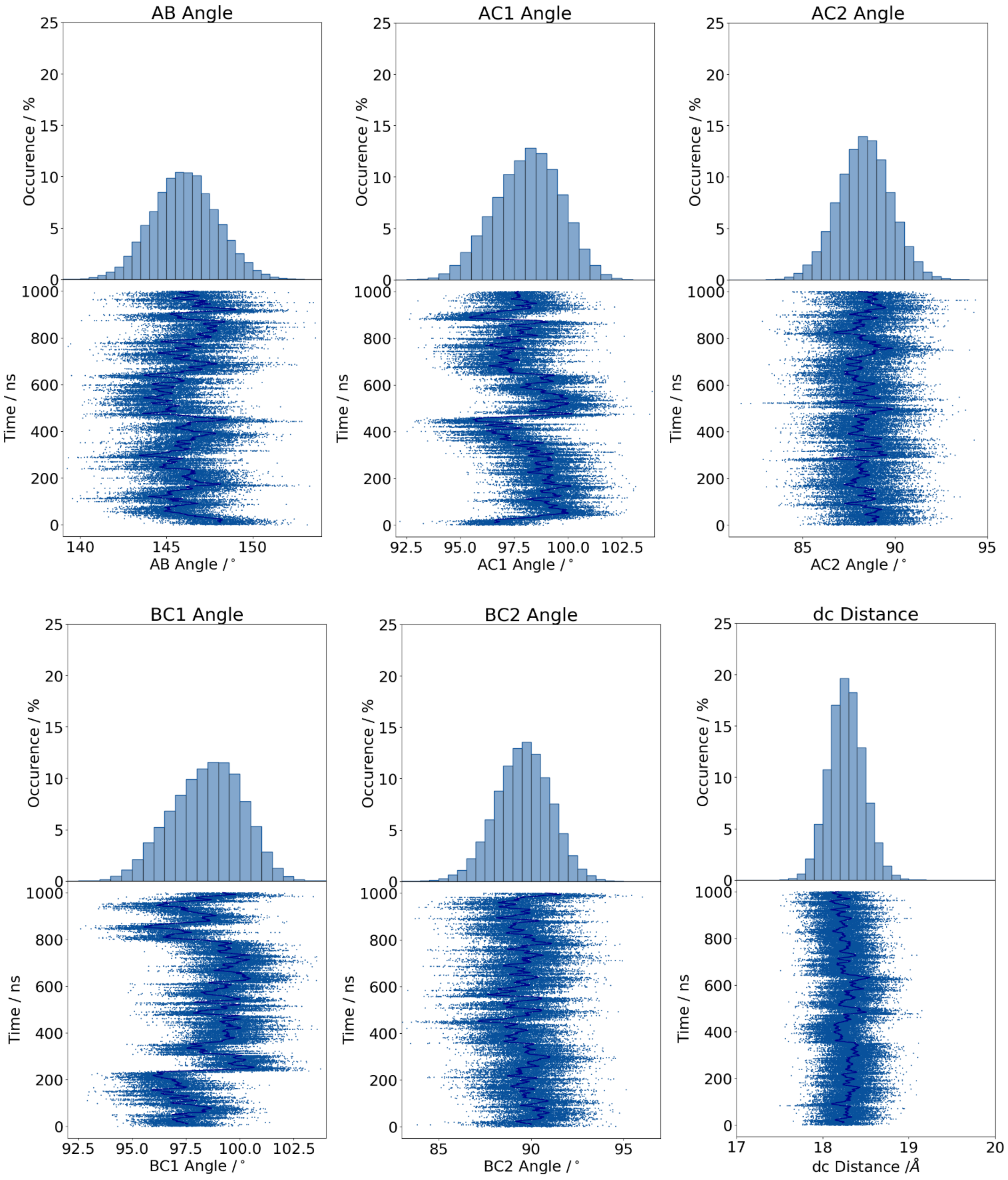


*Figure S 6: OCD Measures of the $C_H3$-$C_H3$ interface. The first frame of the simulation was used as reference structure. Only Cα atoms were included in the calculation. Histograms of the observed data are shown on top, with the actual data in the plot below. To visualize trends in the time series of the measures better, a rolling average over 100 frames is plotted in dark blue on top of the data points.*

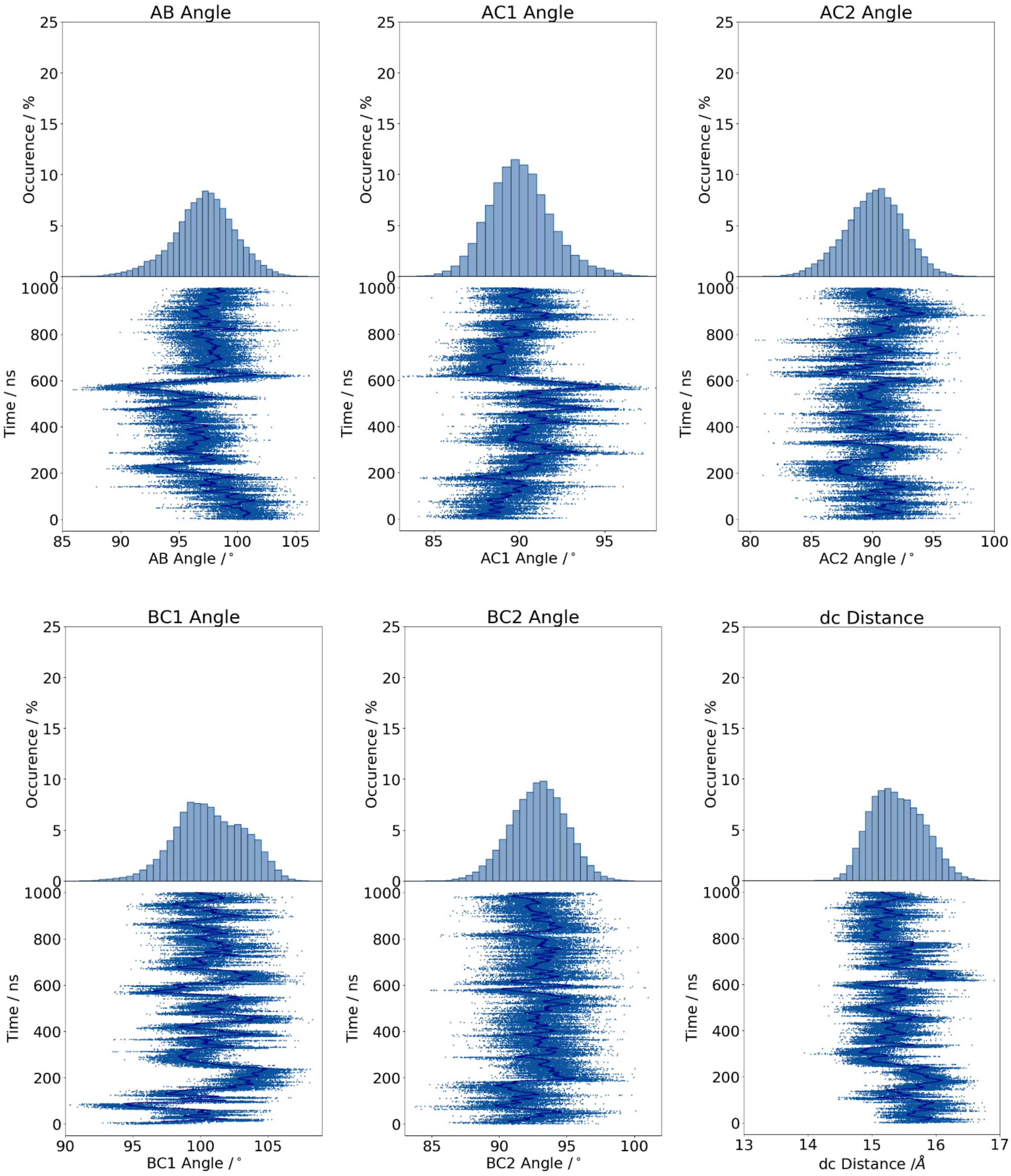


*Figure S 7: OCD Measures of the CH1-CL interface. The first frame of the simulation was used as reference structure. Only Cα atoms were included in the calculation. Histograms of the observed data are shown on top, with the actual data in the plot below. To visualize trends in the time series of the measures better, a rolling average over 100 frames is plotted in dark blue on top of the data points.*
